# Competitive assembly resolves the stoichiometry of essential proteins in infectious HIV-1 virions

**DOI:** 10.1101/2024.03.10.584319

**Authors:** Haley Durden, Benjamin Preece, Rodrigo Gallegos, Ipsita Saha, Brian MacArthur, Abby Petersen, Wiley Peppel, Saveez Saffarian

## Abstract

During assembly on the plasma membrane, HIV-1 virions incorporate Gag-Pol as well as gp120/gp41 trimers. The Pol region consists of protease, reverse transcriptase and integrase precursors which are essential enzymes required for maturation, reverse transcription, and integration of the viral genome in the next host. gp120/gp41 trimers catalyze the fusion of the virion with its next host. Only a fraction of released virions are infectious. The stoichiometry of gp120/gp41 and Gag-Pol proteins in HIV virions was previously measured using cryotomography and ratiometric protein analysis, but what is the stoichiometry of these proteins in infectious virions remained to be determined. Here we developed a method based on competition between infectious HIV backbones with noninfectious mutants and measured 100 ± 10 Gag-Pol and 15 ± 3 gp120/gp41 proteins incorporated in infectious virions assembled in HEK293 cells from NL4.3 HIV-1 backbone. Our measurements are in broad agreement with cryotomography and ratiometric protein analysis and therefore stoichiometry of gp120/gp41 and Gag-Pol in infectious virions is the same as all released virions. With the development of appropriate mutants and infectivity assays, our method is applicable to other infectious viruses.

**Statement of significance:** There are 30 million people who have succumbed to the AIDS pandemic with 600,000 additional deaths per year. HIV has an accelerated rate of mutational accumulation with the virus mutating out of neutralizing antibodies within the same patient making development of vaccines challenging. Like most enveloped viruses, only a fraction of released virions are infectious and the question of what selects these virions has remained a mystery. The method developed in this article will allow stoichiometric measurements on infectious virions and therefore allows further studies of causes of infectivity.

## Introduction

Acquired immunodeficiency syndrome (AIDS) is caused by Human immunodeficiency virus (HIV) which is a major pathogen with a significant death toll around the globe (1). Currently, inhibiting protease, reverse transcriptase and integrase serve as the basis of antiviral therapies for AIDS (2, 3). HIV has a fast replication cycle (1-2 days) and a short lifespan between viral production and infection of the next cell (2-3 days) (4, 5). While a large diverse population of the virus exists in transmitting individuals, productive infections are predominantly established from single founder virion (6–8) suggesting a bottleneck during initial transmission of the virus. While in vitro infectivity assays are far less sophisticated compared to the biology of HIV transmission in vivo, nevertheless, When released virions are assayed in vitro, only 1 in 100 or 1 in 10,000 are found to be infectious (9, 10). There is evidence that stoichiometry of proteins incorporated in different strains of HIV-1 correlates with infectivity (11) and therefore it remains to be determined if infectious virions released from the same variant incorporate a different amount of Gag-Pol or gp120/gp41 trimers to the population.

HIV-1 is a lentivirus from Retriverdea family and carries two copies of a positive sense RNA as its genome. HIV particles assemble on the plasma membrane by incorporating Gag, Gag-Pol, gp120/gp41 trimers and two copies of gRNA among other cofactors (12). The envelope of the virion is decorated with gp120/gp41 protein trimers(13, 14), which upon binding to CD4 and co-receptor CXCR4 or CCR5 catalyze the fusion of the viral envelope with the plasma membrane of the target cell (15–18). Gag-Pol has the precursors for HIV protease, HIV reverse transcriptase and HIV integrase, all enzymes essential for the viral life cycle of HIV. The released virions are immature (19–21) and maturation is catalyzed by HIV-1 protease which performs a specific series proteolytic cuts in Gag and Gag-Pol proteins (22). This results in formation of mature HIV core encompassing two copies of gRNA bound with reverse transcriptase and integrase enzymes (23–25). HIV-1 reverse transcriptase synthesizes a single DNA molecule from two gRNA molecules incorporated within the virion (26), the mature core released in the cytosol after membrane fusion plays an essential role in delivery of the viral DNA genome bound with integrase complexes to the cell nucleus where it integrates into chromosome of the target cell (2, 27, 28). Activation of this embedded genome by transcription leads to initiation of a new viral cycle in the target cell. Because most steps within HIV-1 lifecycle are catalyzed by enzymes with precursors in Gag-Pol and fusion is catalyzed by the gp120/gp41 trimer, it’s reasonable to assume that the number of these proteins incorporated within virions would affect the ability of the virus to go through a full life cycle as described above.

Stoichiometry of HIV-1 virions has been extensively studied. The number of Gag molecules incorporated in HIV-1 virions is estimated by Cryotomography of immature HIV-1 virions to be approximately 2000 copies per virion (29). Its estimated that there are approximately 120 Gag-Pol molecules per virions, although using Cryotomography, Gag-Pol has not been resolved within the isolated immature HIV-1 virions the number of Gag-Pol molecules within the virions is estimated by the frame shifting frequency and the ratio of Gag to Gag-Pol molecules in populations of released virions (29, 30). The number of gp120/gp41 proteins on the surface of the virions has been estimated using cryotomography and shows great variability based on cell types. For virions assembling from NL4.3 backbones in HEK293 cells its estimated that approximately 5 Env trimers are on each virion (11).

HIV-1 research benefits from the development of robust tissue culture tools enabling the study of the viral life cycle in cultured cells. The viruses generated in our study have all been produced using these tools and infectivity is also defined within this context. Specifically in our study we have used the pNL4.3 backbone which encodes a genome derived from multiple wild type circulating group M HIV-1 viruses (31). Transfection of the pNL4.3 backbone DNA into mammalian cells results in release of infectious HIV-1 virions (32). In our study we use the HEK293 cells which support robust production of HIV-1 in culture. To measure infectivity, we used the TZM-bl cells which are derived from Hela cells with addition of CD4 and a tat activated luciferase gene (32). These cells support a linear reading of infectivity from a wide range infectious units (33). Infectivity within the TZM-bl assay is measured as expression of tat activated luciferase in TZM-bl cells after successful infection with HIV virions.

We also benefited from extensive knowledge about abrogating mutations which interfere with protease activation and gp160 proteolysis. Specifically we used NL4.3:Protease(D25N) mutation which abrogates the activity of the HIV protease (34) and the *NL4.3:Env(506SEKS509)* which has a deficient in proteolytic cleavage site in gp160 and virions incorporating only these protein remain non-infectious (35).

In this manuscript we have taken advantage of the fact that multiple viral backbones, in this study pNL4.3 can be transfected into the same cell and virions produced in these cells will assemble from a pool of proteins and gRNA transcribed and translated from these multiple viral backbones. At first, we present a method which measures the number of plasmids which enter and lead to translation of proteins in HEK293 cells. We will show that HEK293 cells receive between 10 to 300 plasmids during each transfection with a distribution best described by three gaussians centered at 40, 80 and 220 pNL4.3 plasmids. We then show that the infectivity of virions collected from a population of HEK293 cells transfected with a constant amount of DNA mix of pNL4.3 and pLN4.3(mutant) shows distinct infectivity curves. In these competitions, the ratio of pNL4.3(mutant) to pNL4.3 varied between 0 to 100. We then show how infectivity of competitions between NL4.3 and NL4.3(SEKS) can be fully modeled using Monte Carlo simulations to deduce that infectious virions contain 15 ± 3 gp120/gp41 proteins. We then show how infectivity of competitions between NL4.3 and NL4.3(D25N) can be fully modeled using Monte Carlo simulations to deduce that infectious virions contain 100 ± 10 Gag-Pol proteins. We then show how infectivity of competitions between NL4.3 and NL4.3(*ΔΨ)*(36) can be fully modeled using Monte Carlo simulations to deduce that infectious virions contain two gRNA molecules. We then show how infectivity of competitions between NL4.3 and NL4.3(Gag(G2A))(34) cannot be fully modeled using Monte Carlo simulations to deduce the number of Gag molecules in the infectious virions.

The closest modeling we could use to describe the NL4.3 and NL4.3(Gag(G2A)) competitions shows that the only released virions that are infectious from these competitions, have two wild type gRNA molecules. We will discuss these results and their implications for the assembly models of HIV.

## Materials and methods

### Competition Experiments: Cell maitanence and transfections for infectivity experiments

Human embryonic kidney (HEK) 293 cell lines were maintained in T-25 flasks using TrypLE Express Enzyme (Gibco) and DMEM with L-Glutamine, 4.5g/L glucose and sodium pyruvate (Corning) supplemented with 10% fetal bovine serum (Gibco). Cells were incubated at 37°C. Once over 90% confluent, the HEK 293 cells were plated onto 10cm tissue culture plates. Experiments were performed in sets of 6 plates, with each plate containing a different ratio of HIV-1 gRNA (pNL4.3) plasmid DNA and a plasmid containing a single-point mutation. Each plate was transfected with 20ng total of plasmid DNA. These transfections were performed using 600ul Opti-MEM Reduced Serum Medium (Gibco) and 40ul of Lipofectamine 2000 (Thermo Fisher Scientific) for each plate. The cells were incubated at 37°C for 36-48hrs.

### Competition Experiments: Harvesting virions and cell extracts for infectivity and western blot assays

To harvest virions for western blot analysis and luciferase assays, cells and supernatants were collected from each 10 cm plate, the contents of each plate was centrifuged at 5,000 RPM (Ample Scientific Champion S-50D centrifuge) for 10min to separate the cells from the supernatant, and then the supernatant was filtered using a 0.22um syringe filter (Celltreat). Before proceeding with further purification of the virions, 1ml of this filtered supernatant was set aside to be used in a luciferase assay. The cell pellets were washed with PBS (Quality Biological) and then resuspended in 100ul of RIPA lysis buffer (Santa Cruz Biotechnology) containing protease inhibitors. The virions were harvested from the supernatant by centrifugation through 1.5ml of 30% sucrose at 10,000 RPM at 4°C for 2hrs (Beckman L8-70M Ultracentrifuge). The resulting pellet of virions was resuspended in 20ul of RIPA lysis buffer containing protease inhibitors.

### Competition Experiments: Luciferase Assay

To test the infectivity of the virions produced from the HEK293 cells, TZM-bl cells were plated onto a 6-well tissue culture plate and incubated with 1ml of supernatant derived from HEK293 cells transfected with proviral DNA. These cells were then incubated at 37°C for 36-48hrs. A britelite plus reporter gene assay system (PerkinElmer) was then used to assess the number of cells that were infected by the virions.

### Competition Experiments: Western Blotting

Cell extracts were centrifuged for 20min at 15,000 RPM (Eppendorf Centrifuge 5424) before being diluted approximately 8X in RIPA lysis buffer. Virions were diluted approximately 2X in the same lysis buffer. Samples were then denatured in 10ul of Lamelli sample buffer (Bio-Rad Laboratories) containing 5% BME and boiled at 95°C for 5min. The samples were loaded into Mini-PROTEAN TGX precast gels (Bio-Rad Laboratories) and their proteins were separated using SDS-PAGE. After completion of electrophoresis, the proteins were transferred onto a PVDF membrane (Millipore). This membrane was then washed with blocking buffer (LI-COR Biosciences) and stained with anti-HIV-1 p24 monoclonal primary antibody (183-H12-5C, NIH HRP) overnight. An anti-mouse secondary antibody (LI-COR Biosciences) was then used to immunoprobe the membrane for 45min before it was scanned with the Odyssey Infrared Imaging System (LI-COR Biosciences) at 700 nm following manufacturer protocol.

### Competition Experiments: Monte Carlo Simulations

To calculate the infectivity of virions released for a range of NL4.3 to NL4.3(mutant) ratios Monte Carlo simulation 10,000 cells were modeled. In each cell, the program assigned a total number of plasmids based on the distribution of number of plasmids from Figure 1. The following ratios of NL4.3/(NL4.3+NL4.3(mutant)) were used (1, 0.9, 0.8, 0.7, 0.6, 0.5, 0.4 0.3, 0.2, 0.1, 0.08 0.07 0.06 0.05 0.04 0.03 0.02 0.01 0). In each cell, based on the assigned total number of plasmids to the cell, number of NL4.3 and NL4.3(Mutant) plasmids were assigned based on the above ratios. The program would run through all 10,000 cells for each one of the ratios presented above. Once the total number of plasmids and the number of the NL4.3 and NL4.3(mutant) plasmids in the cell were assigned, the ratio of mutant and wild type proteins in the cytosol were put proportional to the number of NL4.3 and NL4.3(mutant) plasmids respectively. It was assumed that each cell produces a set number of virions, for simplicity we assumed this number to be 1,000. The number of proteins in each virion was assumed to be “N” and for each virion, the program would randomly select proteins from the mix of wild type and mutant proteins in the cytosol. At the end, the program would have assembled 1,000 virions from each of the 10,000 cells modeled which would make the total number of virions modeled to be 10 million. It is not known how the infectivity of each virion would vary based on the number of mutant versus wild type protein incorporated in the virion. We therefore used a minimum number of wild type proteins required for infectivity as “n”, which is unknown and needs to be fixed when fitting the model to the infectivity data. The Monte Carlo experiments as described above have only two free parameters, N: the total number of proteins of interest in the virion and n: the number of wild type proteins in each virion required for infectivity.

**Figure 1.**
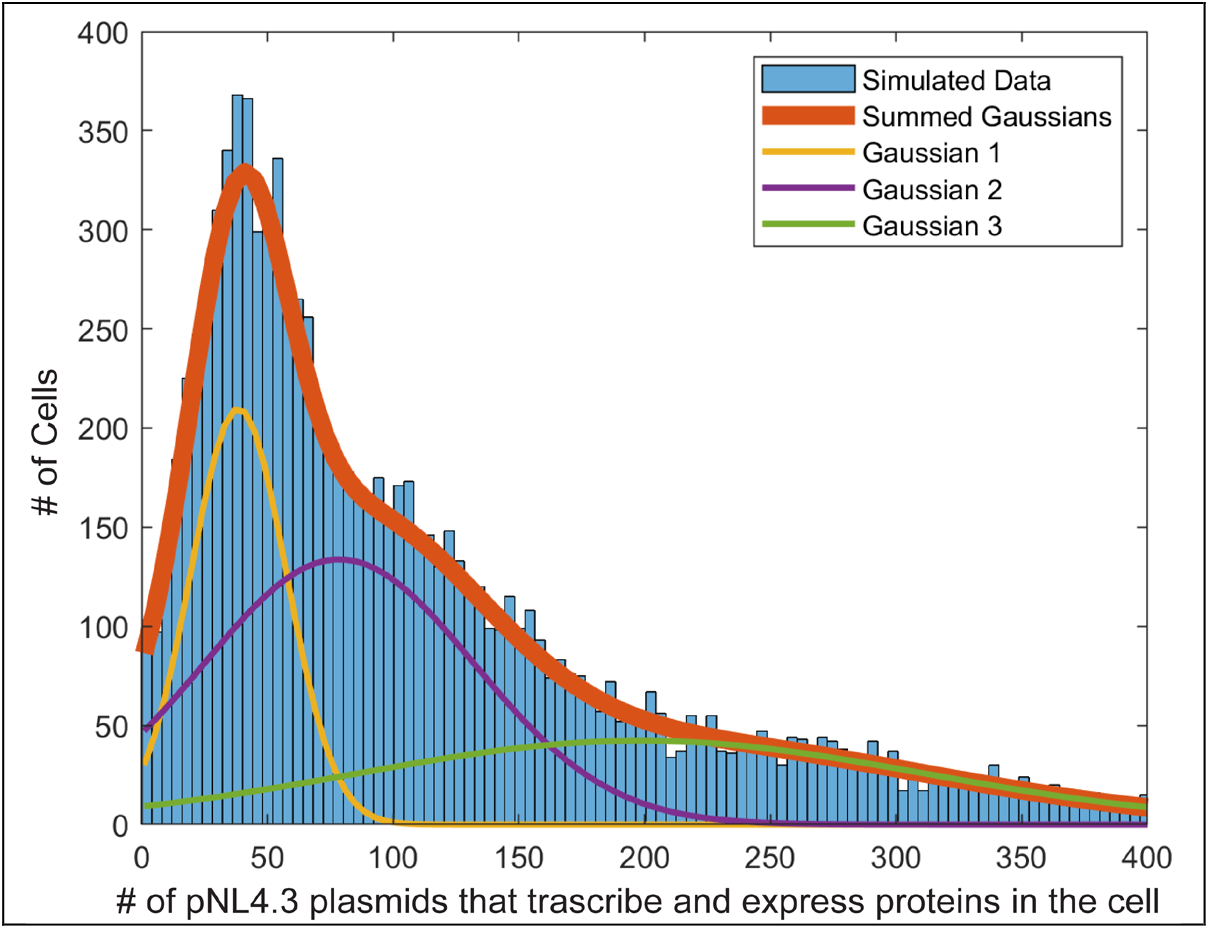
Distribution of the number of pNL4.3 plasmids that enter each cell HEK293 cell after transfection and lead to successful transcription and expression of proteins (N). The distribution does not consider the cells that did not get transfected which are estimated to be around 25% of cells in culture. The distribution of active cells is best described by three gaussians with averages of N = 40 (Gaussian 1 yellow), N = 80 (Gaussian 2 purple) and N = 220 (Gaussian 3).

### Cell plating and transfections for counting the number of plasmids

HEK 293 cells were plated onto 6-well tissue culture dishes with 2 mL of medium per well and incubated for 24 hours prior to transfection. At 50% confluency, the HEK 293 cells were co-transfected with NL4.3(iGFP)(D25N), NL4.3(imCherry)(D25N), and pNL4.3(D25N) at varying dilutions per well. Between the 3 plasmids, 4ng of plasmid (for each well of the 6 well dishes) were used for each transfection at dilutions between 1:2 and 1:256 for the imCherry and iGFP plasmids, with the remainder of the 4ng being pNL4.3(D25N) plasmid. The goal for using the different dilutions was to reach the stochastic limit for random distribution of the plasmids within the cells. Upon reaching said limit, a fraction of cells would only have one type of fluorophore coding plasmid and thereby only express a single-colored fluorophore. Transfection was carried out using 100 ul of Opti-MEM Reduced Serum Medium (Gibco) and 12 ul of Lipofectamine 2000 (Thermo Fisher Scientific) per well. Images were taken 24-48 hours post transfection using an Olympus CKX53 inverted microscope equipped with an Olympus EP50 camera, a brightfield illumination source, and a CoolLED pE-300 light source for red and green fluorescence excitation. All dilutions were imaged using brightfield, 488 nm, and 561 nm light sources to collect background images and individually excite iGFP and imCherry proteins.

## Results

To setup a competition inside the cytosol, where ‘wt’ gRNA and ‘mutant’ gRNA are transcribed and both contribute to the assembly of HIV-1 virions on the plasma membrane of the cell, one needs to first characterize the cells. In our competitions we use HEK293 cells because they are easily transfectable and produce a lot of HIV-1 virions from pNL4.3 backbones. However, the number of backbones which enter each cell after transfection and successfully transcribe and express proteins in each cell is unknown. The distribution of number of plasmids is of great importance for our study since it determines the limits of wt/(wt+mutant) gRNA that can be effectively produced in each cell and it is also important for modeling the competitions from a population of cells.

### Stochastic method for counting the number of plasmids expressing proteins upon transfection into HEK293 cells

To measure the number of plasmids expressing proteins in each HEK293 cell after transfection, we designed our experiments based on detection of fluorescence with the following principles:

If a cell incorporates a number of plasmids “N”, all of equal size and backbone, with a fraction “f, (0 < f < 1)” encoding a red fluorescent protein and a similar fraction “f” incorporating a green fluorescent protein. The fraction of plasmids incorporating no fluorescence can be calculated as:

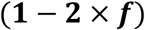

The fraction of fluorescent plasmids (f) can be varied experimentally and can range from 0.5 to extremely small fractions f=1/256. Observing the green and red fluorescent cells in culture upon transfection of various f dilutions can immediately provide an estimate of N. for example, if N=10, which means that each HEK293 cell takes up 10 plasmids, and f = 0.5, the chances of finding a cell that only expresses a red or green color would be ∼(0.5)10 which is 10-3, which means that for every 1000 cells that express fluorescence, only one would have either green or red fluorescence. However, the reality is more complex, with the culture containing a mix of cells, each taking up a different N. Therefore, it’s best to think of N as a distribution.

Figure S1 shows the results of fluorescence imaging in cells transfected with a varying fraction of NL4.3(iGFP), NL4.3(imCherry) and NL4.3 plasmids. For details on the experimental methods for plating and quantifying images, please refer to the section below with the subtitle (Cell plating and experimental details in the stochastic method for counting the number of plasmids).

To numerically calculate the distribution of N within the cell culture, we first quantified the results of experiments presented in Figure S1. Images were analyzed using Fiji imageJ software. Some examples of single-color fluorescence cells are indicated by arrows in figure S1. These results were tabulated using Microsoft Excel. Only cells that expressed imCherry and/or iGFP were counted, while non-fluorescent cells were excluded from calculations. Out of all cells that showed fluorescence, percentages for only eGFP expression, only mCherry expression, and co-expression were tabulated and are presented in Table S1.

A computer simulation was then set up using Matlab to model the observed fluorescence distribution within a system of 10000 cells. The code for this simulation is provided at Github:

(https://github.com/saveez/SaffarianLab/blob/master/HIV-1genomeCompetition.m).

The number of plasmids absorbed per cell was varied until the simulated fluorescence percentages matched our experimental data. By adjusting the number of plasmids expressing in each cell, based on varying distributions, we could explore possibilities for said distribution until we found a close fit for our experimental data. Ultimately, the plasmid uptake distribution was concluded to be best modeled by a set of three gaussian curves as shown in Figure 1. This places the overall average around 100 plasmids per cell, with most cells expressing between 10 to 300 plasmids each.

### Competitions between NL4.3 and NL4.3(protease(D25N)) “inactive Protease”

To test the assembly of infectious HIV in the presence of a mix of NL4.3 and NL4.3:Protease(D25N) plasmids competition experiments were set up as described in methods and briefly summarized here:10 cm plates of HEK293 cells were transfected with a constant amount of DNA with varying ration of NL4.3:Protease(D25N) to NL4.3 from zero to hundred. Virions released from these cells were incubated with TZM-bl cells and infectivity was read by luciferase activity within these cells. Western blots were performed on cells and released virions to verify protein expression and levels of proteolysis shown in Figure 2.

**Figure 2.**
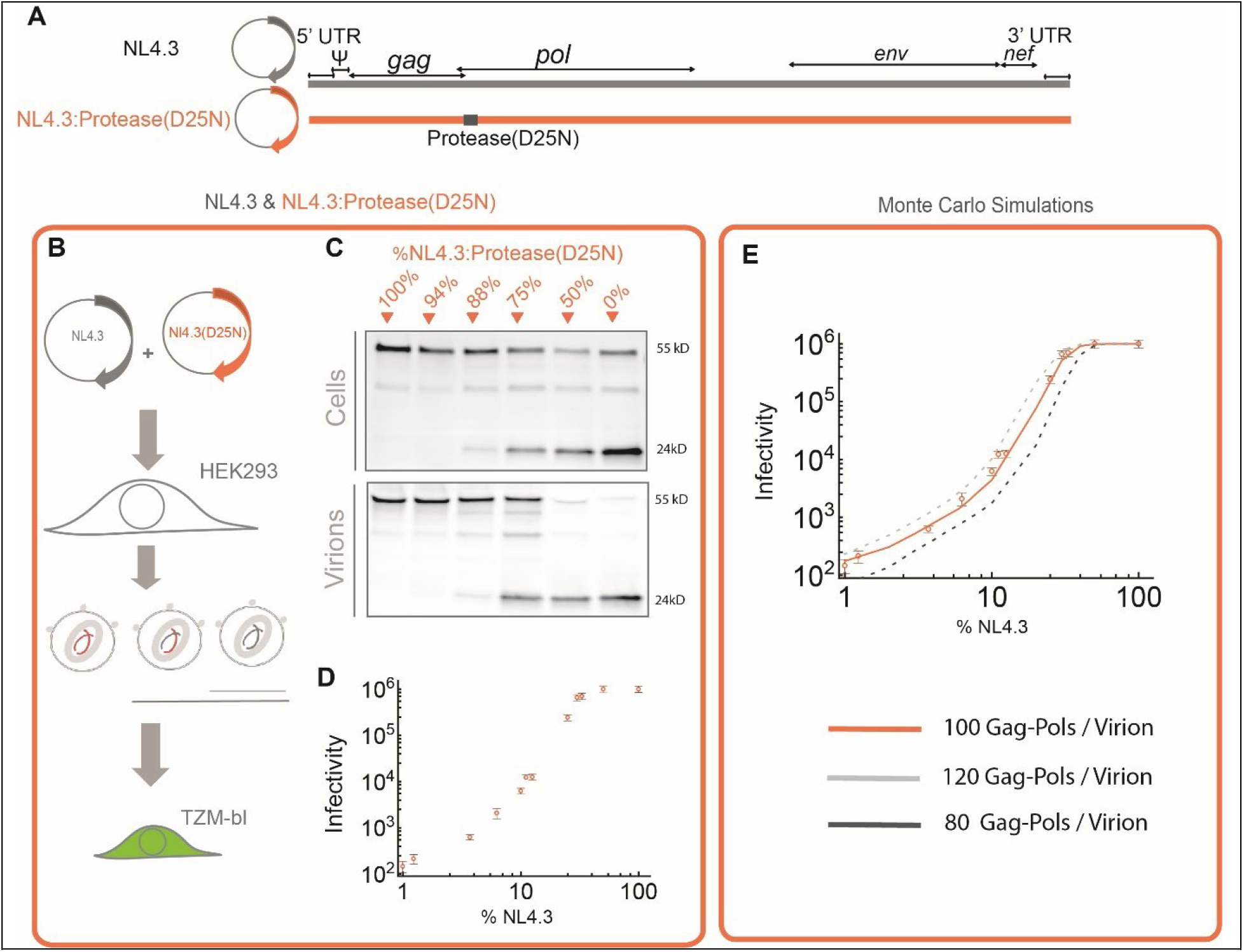
HIV-1 assembly competitions between NL4.3 and NL4.3:Protease(D25N). The genome maps for NL4.3, NL4.3:Protease(D25N) (A). Schematic diagram of the competition experiments (B), Virions released by co-assembly of NL4.3 and NL4.3:Protease(D25N) in HEK293 cells assayed with western blots (C) and infectivity in TZM-bl cells (D). Monte Carlo simulations based on assembly of the virions on the plasma membrane and random incorporation of Gag-Pols within assembling virions on the plasma membrane from the cytosol (E).

In competitions between NL4.3 and NL4.3:Protease(D25N), the amount of released virions, as visualized in western blots, is constant regardless of the ratio of NL4.3 to NL4.3:Protease(D25N) (Figure 2.C). As the fraction of NL4.3 is decreased, the infectivity of virions initially remains constant and starts to decline when about 30% of DNA is from NL4.3 and correspondingly 70% is from NL4.3:Protease(D25N). This decline continues with lower fractions of NL4.3, however the infectivity of virions never reaches zero even when only 1% of the DNA is from NL4.3 and correspondingly 99% is from NL4.3:Protease(D25N).

### Analysis of the NL4.3:NL4.3(protease(D25N)) competition using Monte Carlo simulations

In our simulations we assumed that during the competition experiments, each cell received a mix of NL4.3 and NL4.3(protease(D25N)) plasmids. Based on the current model of assembly, these plasmids would get transcribed and generate a set of NL4.3 and NL4.3(protease(D25N)) gRNA in the cytosol. Gag and Gag-Pol would get translated from these gRNA and accumulate within the cytosol. Once assembly on the plasma membrane is initiated, the assembling virion would be populated by the Gag and Gag-Pol molecules translated from the pool of gRNA. The Monte Carlo simulation described in the methods and shown in Figure S2, models the assembly of 10 million virions from 10,000 cells each incorporating a set of NL4.3 and NL4.3(protease(D25N)) plasmids based on distribution shown in Figure 1 and determines the fraction of virions which are infectious.

The model has only two parameters, N: which is the total number of Gag-Pol molecules assembling in each virion and n: which is the number of wild type Gag-Pol molecules needed for the virion to be infectious. Curves generated by this model mimic the shape of the experimental infectivity data. Best fits are achieved when n=30. The total number of gag-Pols N can be varied and results in horizontal shifts in the infectivity curves. Figure 2 shows results of the Monte Carlo simulations with N=80, N=100 and N=120. The best fit is achieved with N=100. The spread of data with the fit dictates an error of ±10 in our measurements of N, therefore we conclude that infectious virions incorporate 100 ± 10 Gag-Pol molecules.

### Competitions between NL4.3 and NL4.3:Env(506SEKS509) “gp160 resistant to proteolysis”

To test the assembly of infectious HIV in the presence of a mix of NL4.3 and NL4.3:Env(506SEKS509) plasmids competition experiments were set up as described in methods and briefly summarized here:10 cm plates of HEK293 cells were transfected with a constant amount of DNA with varying ration of NL4.3:Env(506SEKS509) to NL4.3 from zero to hundred. Virions released from these cells were incubated with TZM-bl cells and infectivity was read by luciferase activity within these cells. Western blots were performed on cells and released virions to verify protein expression and levels of proteolysis shown in Figure 3.

**Figure 3.**
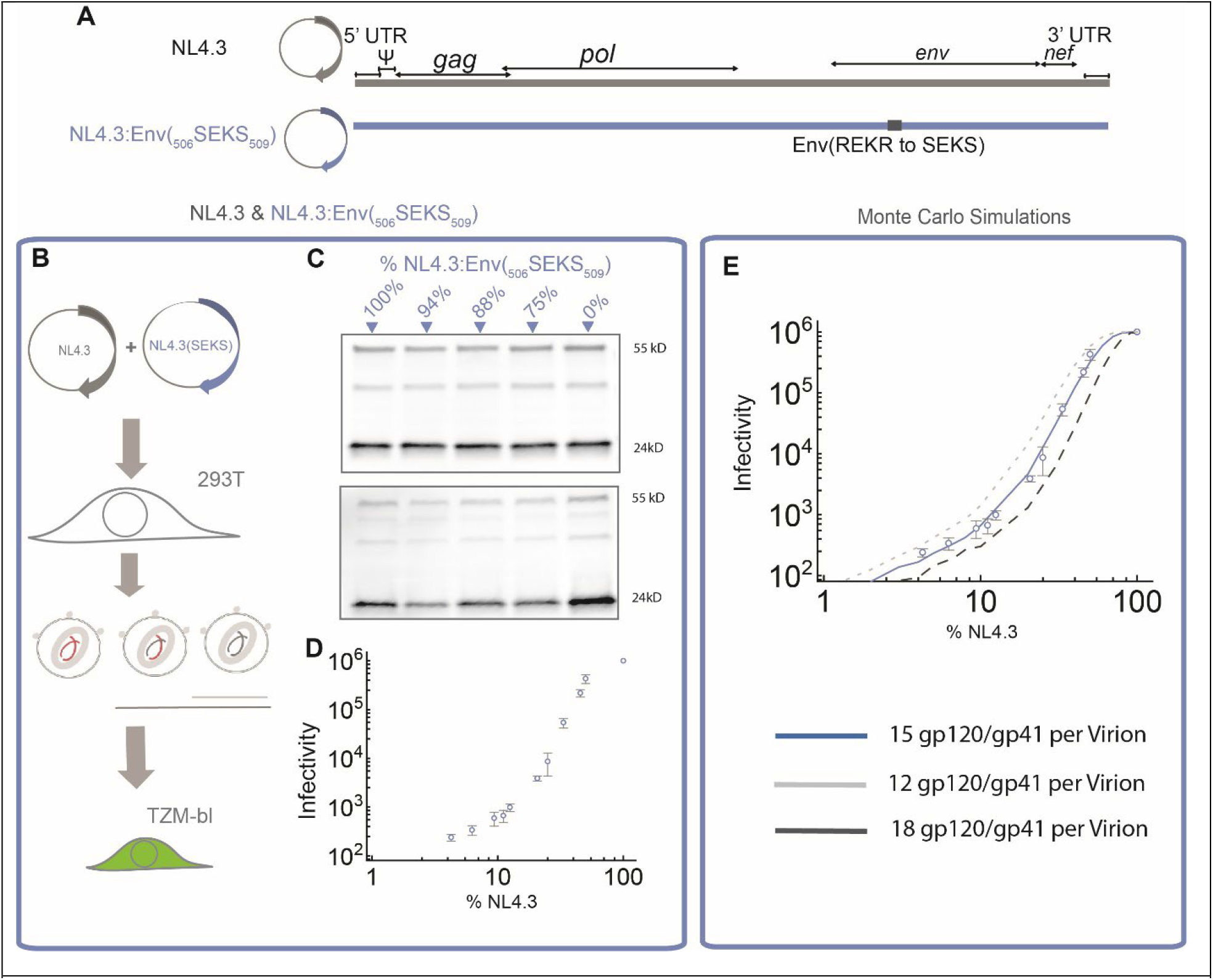
HIV-1 assembly competitions between NL4.3 and NL4.3:Env(_506_SEKS_509_). The genome maps for NL4.3, NL4.3:Env(_506_SEKS_509_) (A). Schematic diagram of the competition experiments (B), Virions released by co-assembly of NL4.3 and NL4.3:Env(_506_SEKS_509_) in HEK293 cells assayed with western blots (C) and infectivity in TZM-bl cells (D). Monte Carlo simulations based on assembly of the virions on the plasma membrane and random incorporation of gp120/gp41 within assembling virions on the plasma membrane from the membrane (E).

In competitions between NL4.3 and NL4.3:Env(506SEKS509), the amount of released virions, as visualized in western blots, is constant regardless of the ratio of NL4.3 to NL4.3:Env(506SEKS509) (Figure 3.C). As the fraction of NL4.3 is decreased, the infectivity of virions declines. This decline continues with lower fractions of NL4.3.

### Analysis of the NL4.3: NL4.3:Env(_506_SEKS_509_) competition using Monte Carlo simulations

In our simulations we assumed that during the competition experiments, each cell received a mix of NL4.3 and NL4.3:Env(_506_SEKS_509_) plasmids. Based on the current model of assembly, these plasmids would get transcribed and generate splice variants encoding ENV from a set of NL4.3 and NL4.3:Env(_506_SEKS_509_). Gag and gp160 proteins would get translated from these splice variants and make their way to the plasma membrane. We assume that the gp160 translated from NL4.3 would undergo proteolysis and generate gp120/gp41 proteins whereas the gp160 translated from NL4.3:Env(_506_SEKS_509_) would not be proteolyzed, however would traffic and assemble on the virions in a similar fashion as gp120/gp41. Once assembly on the plasma membrane is initiated, the assembling virion would be populated by the gp120/gp41 and gp160 molecules translated from the pool of splice variants in the cell. The Monte Carlo simulation described in the methods and shown in Figure S3, models the assembly of 10 million virions from 10,000 cells each incorporating a set of NL4.3 and NL4.3:Env(_506_SEKS_509_) plasmids based on distribution shown in Figure 1 and determines the fraction of virions which are infectious. The model has only two parameters, N: which is the total number of gp120/gp41 and gp160 molecules assembling in each virion and n: which is the number of gp120/gp41 molecules needed for the virion to be infectious. Curves generated by this model mimic the shape of the experimental infectivity data. Best fits are achieved when n=9. The total number of gp120/gp41 and gp160, N can be varied and results in horizontal shifts in the infectivity curves. Figure 3 shows results of the Monte Carlo simulations with N=12, N=15 and N=18. The best fit is achieved with N=15. The spread of data with the fit dictates an error of ±3 in our measurements of N, therefore we conclude that infectious virions incorporate 15 ± 3 gp120/gp41 and gp160. We assume that in absence of competition, virions assembled from NL4.3 would incorporate 15 ± 3 gp120/gp41 molecules which would be 5 trimers on the envelope of the virions.

### Competitions between NL4.3 and NL4.3:Gag(G2A), NL4.3:ΔΨ(Δ105-278/Δ301-332) “Gag that doesn’t bind membrane, and gRNA with no Ψ packaging signal “

The infectivity data presented in Figures 2&3 was easily modeled using Monte Carlo simulations based on current models of assembly and revealed interesting measurements about the stoichiometry of the infectious HIV virions. In this section, we decided to push these competition further into elements of HIV which are not enzymes and are more structural parts of the virions. For this we decided to test Gag and gRNA in competitions. The NL4.3(ΔΨ(Δ105-278/Δ301-332)) which deletes the complete Ψ packaging signal from the gRNA (36) when transfected into HEK293 cells releases virions that are not infectious when tested in our TZM-bl assay. The G2A mutation within the Gag protein, abrogates the interaction of Gag with the plasma membrane and abrogates release of virions all together (34). It is important to note that both of these mutations/deletions have a structural effect on HIV virions and therefore the simple assumption we had in our Monte Carlo model, in which the recruitment of proteins/gRNA into the assembling virions was blind to the presence of the mutation would not necessarily apply to the above competitions.

The experimental infectivity curves derived from competitions between NL4.3 and NL4.3(ΔΨ(Δ105-278/Δ301-332)) are shown in Figure 4 B and experimental infectivity curves derived from competitions between NL4.3 and NL4.3:Gag(G2A) are shown in Figure 4 C.

**Figure 4.**
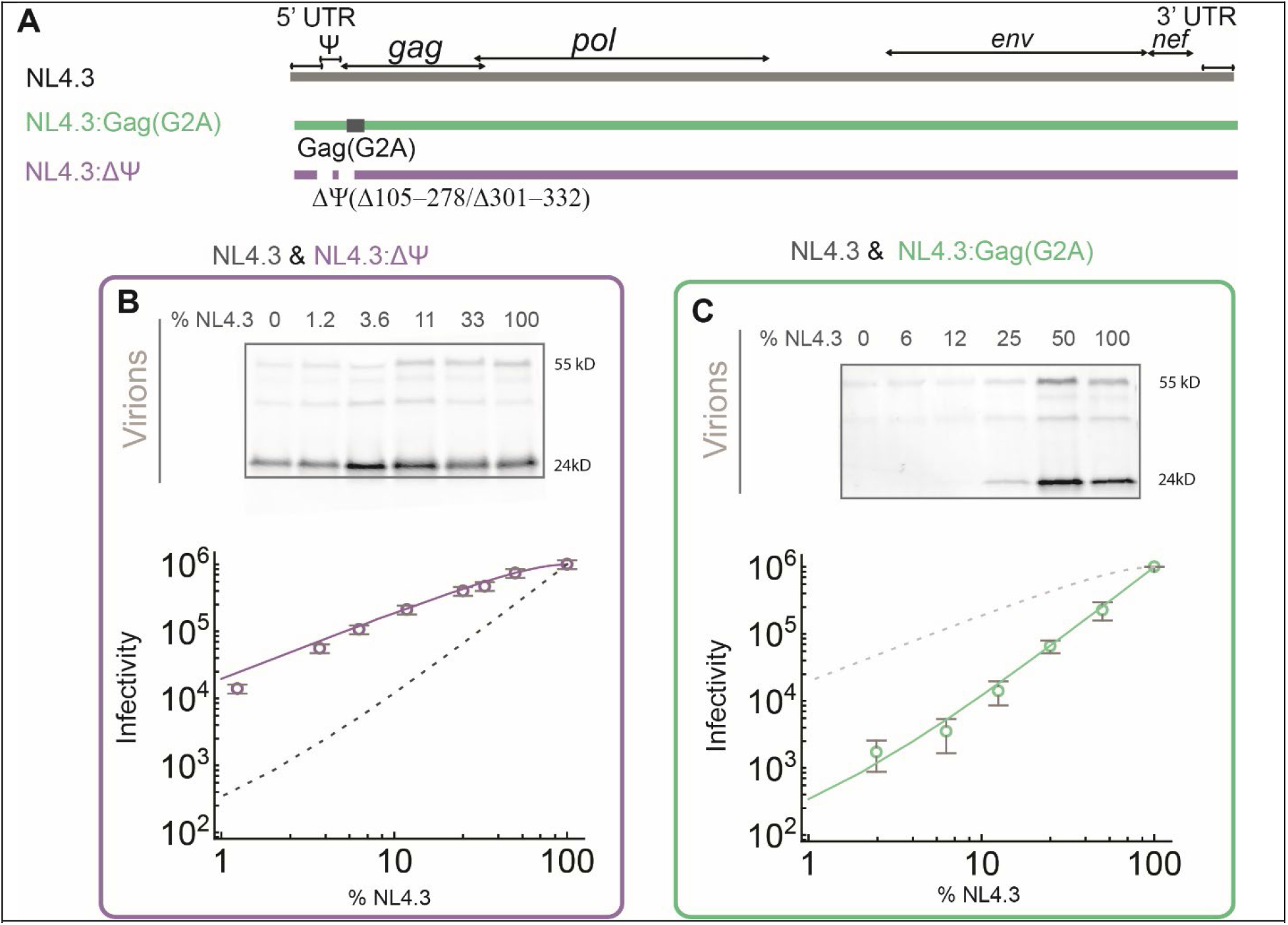
HIV-1 assembly competitions between NL4.3 vs NL4.3:ΔΨ(Δ105-278/Δ301-332) and NL4.3 vs NL4.3:Gag(G2A). The genome maps for NL4.3:ΔΨ(Δ105-278/Δ301-332) and NL4.3:Gag(G2A) (A). Competition between NL4.3 and NL4.3:ΔΨ(Δ105-278/Δ301-332) with western blots showing the release of virions at different percentages of NL4.3 and infectivity data modeled by a Monte Carlo simulation showing that two gRNA’s are incorporating in every released infectious virion with at least one wild type gRNA required for infectivity (B). Competition between NL4.3 and NL4.3:Gag(G2A) with western blots showing the release of virions at different percentages of NL4.3 and infectivity data modeled by a Monte Carlo simulation showing that the release of infectious virions in this competition is proportional to the fraction of virions which incorporate two wild type gRNAs.

### Analysis of the NL4.3 vs NL4.3:Gag(G2A) and NL4.3 vs NL4.3:ΔΨ(Δ105-278/Δ301-332) competition using Monte Carlo simulations

In our simulations we assumed that during the competition experiments between NL4.3 vs NL4.3:ΔΨ(Δ105-278/Δ301-332), each cell received a mix of NL4.3 and NL4.3:ΔΨ(Δ105-278/Δ301-332) plasmids. Based on the current model of assembly, these plasmids would get transcribed and generate a pool of gRNA in the cytosol which had both gRNA transcribed from NL4.3 and gRNA transcribed from NL4.3:ΔΨ(Δ105-278/Δ301-332), we assumed that the population of these gRNA’s reflected the population of plasmids which entered the cells. We assumed that both gRNA species would translate Gag, Gag-Pol and all other HIV co-factors. Once assembly on the plasma membrane is initiated, we assumed that the virions would have N number of gRNAs from which n had to be wild type gRNA. The data as presented in Figure 4B shows that the best fit is achieved when N=2 and n=1, which means that each infectious virion has to incorporate two gRNAs and at least one of these gRNA species has to have an intact Ψ packaging signal.

In our simulations we assumed that during the competition experiments between NL4.3 vs NL4.3:Gag(G2A), each cell received a mix of NL4.3 and NL4.3:Gag(G2A) plasmids. Based on the current model of assembly, these plasmids would get transcribed and generate a pool of gRNA in the cytosol which had both gRNA transcribed from NL4.3 and gRNA transcribed from NL4.3:Gag(G2A), we assumed that the population of these gRNA’s reflected the population of plasmids which entered the cells. We assumed that both gRNA species would translate Gag, Gag-Pol and all other HIV co-factors. And therefore the population of Gag(G2A) in the cytosol of each cell, would be proportional to the G2A gRNA present in the cytoplasm. The G2A competitions have a non-trivial infectivity curve. We first assumed that each virion has to incorporate N=2000 gag molecules and only a fraction of these needed to be wild type to rescue infectivity of the virions. These simulations which are shown in Figure S4 did not match the experimental data. We then noticed that if we plot the experimental data from the G2A competition, it overlaps nicely with the expected infectivity from virions that incorporate two wild type gRNAs. We present this line in Figure 4C, while we do not have a very clear explanation for this fit and would be discussing this further in our discussion.

## Discussion

HIV-1 is a lentivirus and understanding its assembly has been fundamental in development of lentiviral vectors (37, 38). Only a small fraction of released HIV-1 virions are infectious, based on data presented in our manuscript, for HIV-1 virions generated from HEK293 cells by transfection of pNL4.3 backbone, we find that less than 1% of virions are infectious. As outlined in the introduction, what selects the infectious virions remains poorly understood. Mathematical modeling has been previously used to link the number of gp120/gp41 proteins found in different variants with infectivity of the virus(9, 11, 39). However natural variations of the HIV proteins found within a population of virions all generated from the same backbone has been more challenging to explore.

The competition experiments introduced in this manuscript utilize a combination of modeling with quantitative infectivity assay to deduce the composition of the infectious HIV virions. As explained in the introduction section, the infectivity in our manuscript is strictly defined as virions that can infect TZM-bl cells and induce the luciferase gene in these cells. This particular definition does not include the passage of the virus beyond its first host, something that many of the infectious virions defined in our study would not be able to do, because they carry protease mutations or ENV mutations in their gRNA and therefore would have difficulty replicating. Because the mutations within the gRNA molecule in the protease gene and ENV gene do not have any structural effects on the viral particles, the infectivity is only reflecting protein effects and allows for the competition to be modeled easily.

As for quantification of stoichiometry of gp120/gp41 proteins, our measurements are broadly consistent with previous estimates for NL4.3 backbones assembling in HEK293 cells which found approximately 5 Env trimers are on each virion (11). Since the previous measurements were performed using CryoEM, they reflect an average of 5 trimers within the population of virions. We conclude from these comparisons that the infectious and non-infectious virions released from the HEK293 cells after transfection with the pNL4.3 backbone share similar stoichiometry of gp120/gp41 proteins and unlike observed differences between variants of HIV which resulted in differences in infectivity of virions (9, 11, 39), within the same variant, infectious and non-infectious virions have the same stoichiometry.

The model used to fit the data in our study is consistent with the current understanding of HIV assembly where the viral gRNA serves as a template for ribosomal translation of Gag and Gag-Pol while the gp160 the precursor of gp120/gp41 is translated from a splice variant (40–42). Gag-Pol is translated after a (−1) ribosomal slippage with 10-20 Gag-Pol molecules for every 100 Gag molecule translated which is likely regulated by t-RNA levels (30, 43, 44). Binding of Gag to membrane, gRNA and tRNA (34, 36, 45–47), loading of Gag-Pol within virions (30, 48), recruitment of gRNA and its dimerization (49–54) and the choreography of HIV-1 protease activation (22) has been extensively investigated. Our quantification of stoichiometry of Gag-Pol within infectious virions is in broad agreement with the stoichiometry of Gag-Pol estimated by combining the frameshifting frequency and the estimated number of Gag molecules found in purified immature virions(29).

Therefore, we did not find evidence to support the hypothesis in which infectious virions would incorporate more or less of the Gag-Pol protein.

The competition experiments between NL4.3 and NL4.3:ΔΨ(Δ105-278/Δ301-332) show that the infectious virions do incorporate two viral genomes which is consistent with previous observations of gRNA recruitment and its dimerization (49–54). The fact that our models predict that virions which incorporate one wild type gRNA can also include one gRNA with ΔΨ(Δ105-278/Δ301-332) is interesting, however we have not verified these predictions by analyzing the gRNA molecules in the released virions and therefore while the math and model supports this claim, this analysis does not take into account the role for the Ψ packaging signal during assembly and therefore can be misleading.

Similar to the NL4.3 and NL4.3:ΔΨ(Δ105-278/Δ301-332) competitions, the competition between NL4.3 vs NL4.3:Gag(G2A) is not easily modeled because the Gag protein plays a pivotal role during HIV assembly and one cannot assume that Gag and Gag(G2A) molecules would be recruited into assembling virions with the same efficiency. We believe that this is the reason why the Monte Carlo simulation with such an assumption as presented in Figure S4 can not fit the observed infectivity in the NL4.3 vs NL4.3:Gag(G2A) competition. It is however curious that the infectivity trend in NL4.3 vs NL4.3:Gag(G2A) tracks the fraction of virions which would statistically incorporate two wild type genomes. Similar to the disclaimer about ΔΨ, the mathematical prediction that infectious virions would have two wild type genomes has not been verified by additional experiments in our manuscript, however if one could setup a system where infectious and non-infectious virions could be separated and structurally analyzed, the mathematical claim could be verified.

We conclude by stating that HIV assembly is complex and further studies examining the causes of infectivity are warranted.

## Author Contributions

HD, BP, BM, AP, WP, RG performed experiments and quantified data. IS and SS designed experiments, SS wrote the manuscript.

## Acknowledgments

Funding: This research was funded by NIH R56 AI150474-06A1 to (SS)

