## Supplemental figures and text for "Competitive assembly resolves the stoichiometry of essential proteins in infectious HIV-1 virions"

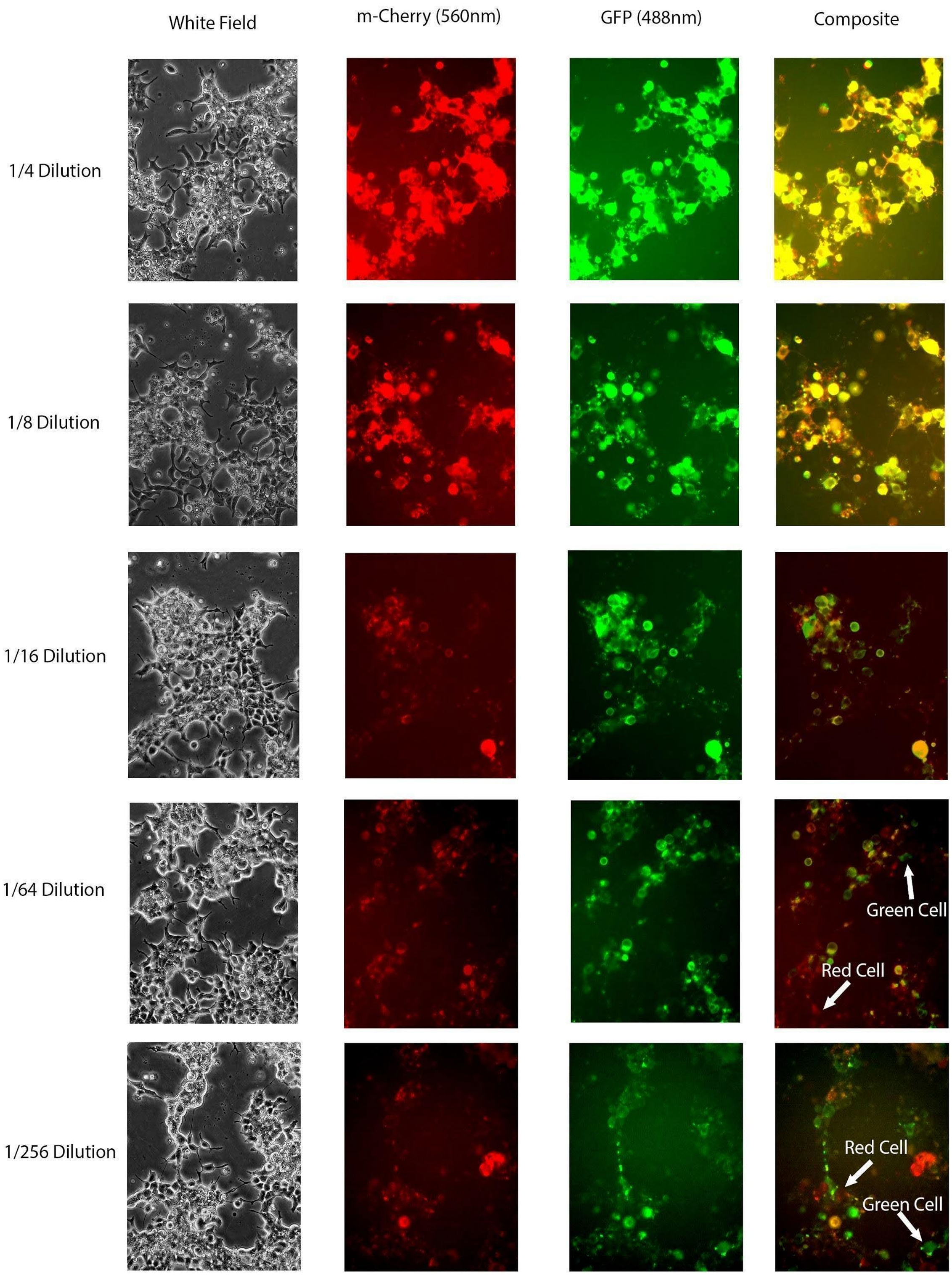


**Figure S1.** Images from Cell Dilution sets. Examples include brightfield, blue illuminated (iGFP excitation), green illuminated (imCherry excitation), and green/blue composite images taken. Dilutions on the left indicate the transfection ratio of iGFP and imCherry coding plasmids to dark plasmids. Examples of single-color cells are highlighted in the last two dilution sets with white arrows.

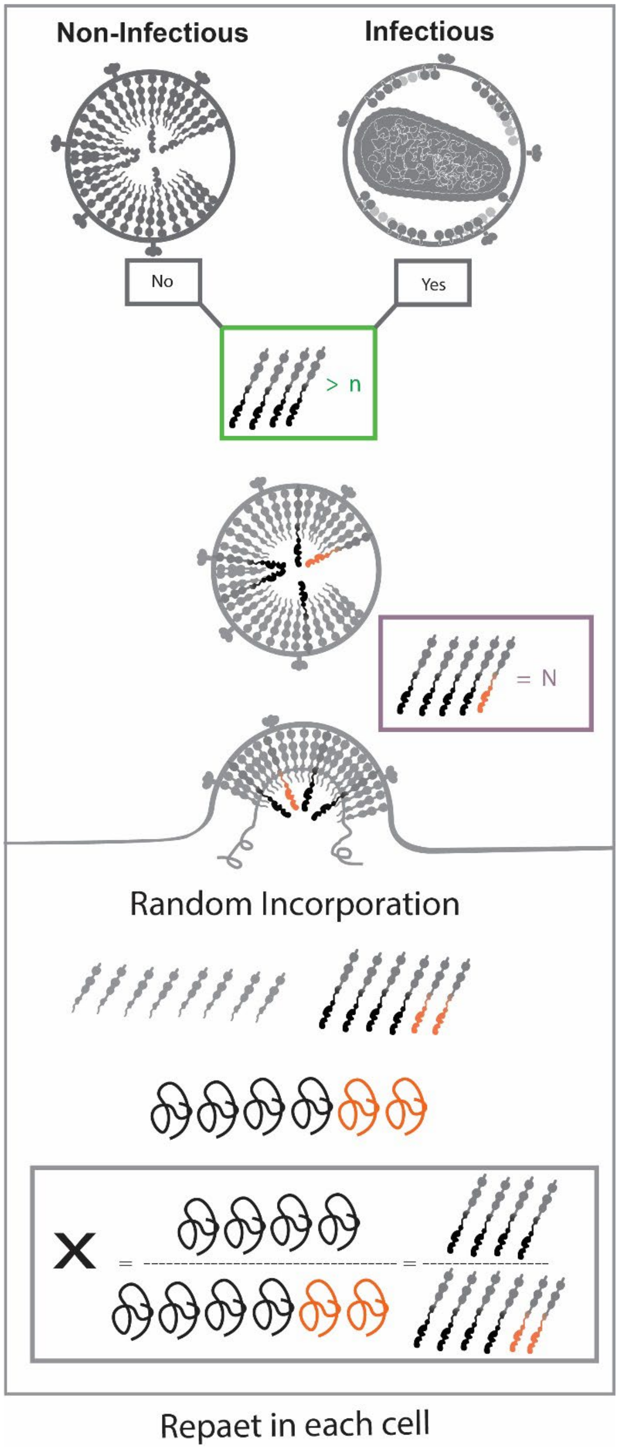


Figure S2: Monte Carlo simulations diagram for NL4.3 vs NL4.3:Protease(D25N) competitions.

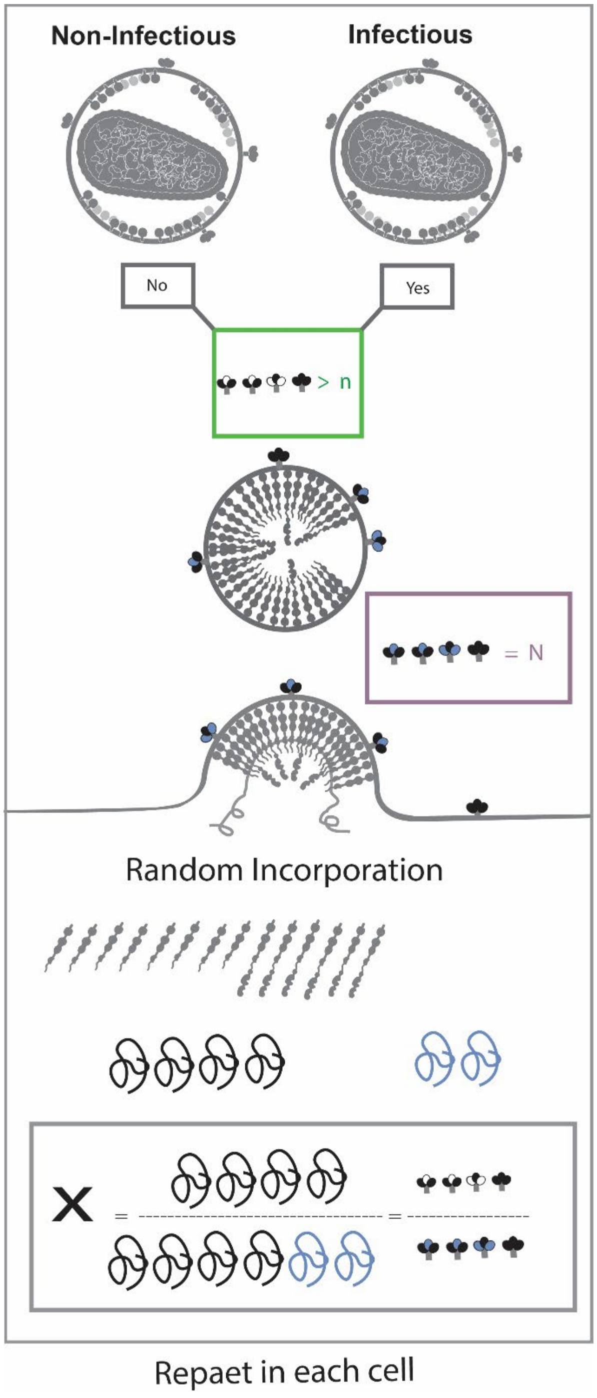


Figure S3: Monte Carlo simulations diagram for NL4.3 and NL4.3:Env($_{506}$SEKS$_{509}$) competitions.

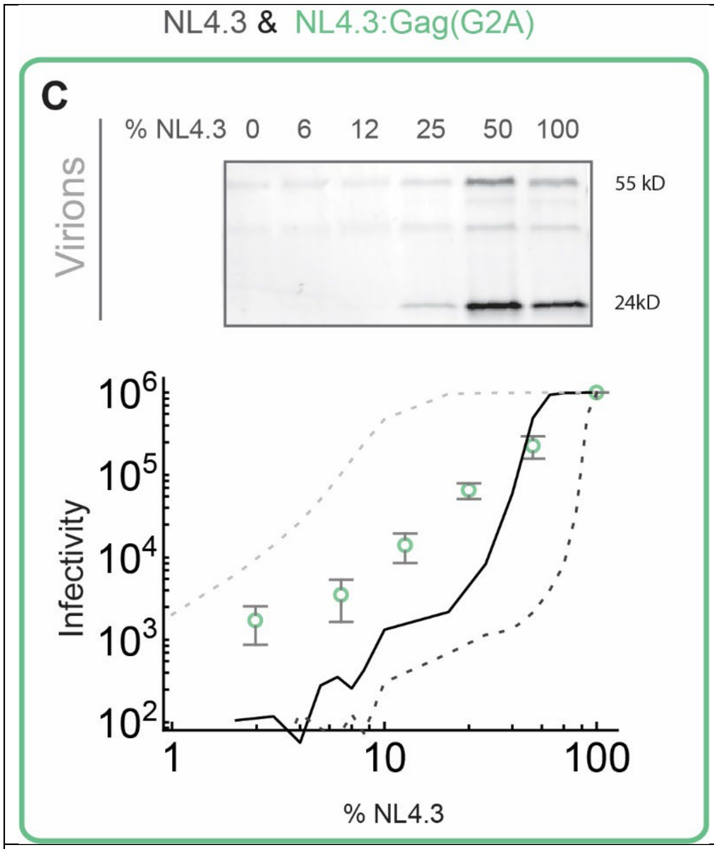


Figure S4. HIV-1 assembly competitions between NL4.3 vs NL4.3:Gag(G2A). Monte Carlo simulations based on 200 Gag molecules N = 2000 with n = 200 (dark dash), n=1000 (Dark line) and n=1800 (Light dash) shown.

| Dilution (f) | Fraction of cells with Green or red only | Both red and green |
|---|---|---|
| 0.00391 | 0.3337 | 0.3457 |
| 0.015625 | 0.1979 | 0.6041 |
| 0.0625 | 0.0584 | 0.8831 |
| 0.125 | 0.0134 | 0.9732 |

**Table S1.**

Fraction of cells expressing either one color or both color in cultures of HEK293 cells transfected with a mix of NL4.3(iGFP), NL4.3(imCherry) and NL4.3(D25N).
